# FASTERCC: Accelerating Flux Consistency Testing and Context-Specific Reconstruction for Large-Scale Metabolic Network Models

**DOI:** 10.64898/2026.03.19.712885

**Authors:** Maria Pires Pacheco, Evelyn Gonzalez, Thomas Sauter

## Abstract

The increase in size of metabolic network models especially with the advent of single-cell data calls for scalable reconstruction and analysis tools. Such models, often used for drug discovery and the analysis of microbial communities rely on consistency testing and reconstruction algorithms such as FASTCORE and FASTCC. However, with models nowadays comprising hundreds of thousands of reactions, the running times of such algorithms increased from few minutes to hours or days even with high performance computing. Experiments that require multiple reconstructions, such as parameter tuning or cross-validation, are practically infeasible in very large networks.

Here we introduce FASTERCC, a new version of FASTCC, that leverages structural information for removing type I and II dead-ends, the orientation of reversible reactions and correcting the reversibility of reactions that are structurally incapable of carrying flux in both directions prior to any feasibility tests. These improvements reduce drastically the running time of FASTERCC by a median 20-fold speedup in comparison to FASTCC for networks with a larger number of block reactions. The model cleaning performed by FASTERCC also reduces the computational time of downstream analyses, notably of FASTCORE up to 50%.

## Introduction

High-throughput technologies, particularly single-cell sequencing, are generating biological datasets of unprecedented scale, calling for efficient and scalable computational analysis workflows. Genome-scale metabolic models (GEMs) have been widely used for predicting repurposable metabolic drugs (Bintener et al., 2023; Lim et al., 2025; Pacheco et al., 2019; Turanli et al., 2019), integrating omics data (Jamialahmadi et al., 2019; Pacheco et al., 2015a, 2022, 2026; Sen & Orešič, 2023; Zagare et al., 2025; Zhang et al., 2019) and microbiome engineering (Quinn-Bohmann et al., 2025) among others. These applications rely on context-specific reconstruction and flux consistency testing algorithms. Context-specific reconstruction algorithms tailor generic reconstructions to specific conditions. Flux-consistency testing algorithms identify reactions that cannot support any steady-state flux (blocked reactions).

FASTCORE and FASTCC (Vlassis et al., 2014) were the first algorithms to achieve these tasks with running times below one minute for genome-scale metabolic networks. Due to their efficiency, they were widely adopted by the community for high-throughput model building and flux consistency testing. However, metabolic models derived from single-cell data or microbial communities can comprise hundreds of thousands of reactions, which challenges existing reconstruction and consistency-testing pipelines. This high number of reactions has made model building and flux consistency testing increasingly computationally expensive, with running times up to several days and, in some cases, infeasible without high-performance computing (HPC) (Varrette et al., 2014). This loss of scalability cannot be solved by more powerful hardware alone but requires more efficient implementations of algorithms. Many analyses requiring multiple reconstructions, such as parameter tuning and cross-validation, are practically impossible even on HPC-based solutions.

A major driver of this computational inflation lies in the handling of reversible reactions and particularly blocked ones; the number of required linear programming (LP) solves grows, becoming the dominant bottleneck in both FASTCC and FASTCORE (Vlassis et al., 2014). While the flux-consistency of all irreversible reactions is assessed simultaneously in a single optimization step, reversible reactions might require multiple feasibility checks due to their orientation. During consistency testing, flux is pushed in the forward direction; hence, a contradictory orientation might prevent flux thereby triggering flipping and retesting of the reversible reactions. The presence of blocked reactions culminates in the costly one-by-one feasibility checks, in which every reversible reaction that did not pass the previous flux feasibility testing is checked individually.

To overcome these scalability issues, we introduce FASTERCC, a structurally informed extension of FASTCC that dramatically accelerates flux consistency testing by leveraging network topology. FASTERCC introduces three key preprocessing steps: (1) correcting the orientation of mis-specified irreversible and reversible reactions, (2) converting structurally constrained reversible reactions into irreversible ones, and (3) detecting and pruning dead-end metabolites (both type I and II) before optimization. By eliminating blocked reversible reactions and reducing the total number of reversible reactions upfront, FASTERCC minimizes the need for costly one-by-one LP solves, achieving a speedup of up to 30-fold for larger number of block reactions (*e.g.*, removal of 500 reactions). Importantly, these structural modifications also improve the performance of downstream tools like FASTCORE, reducing its runtime by up to 50% when FASTERCC is used as a preprocessing step.

## Methods

### 1. FASTERCC algorithm

FASTERCC accelerates flux consistency testing by including a structure-based preprocessing step comprising, the reorientation of irreversible reactions with negative lower bounds, the removal of dead-ends (Figure 1a) and structural tightening of reaction directionality (Figure 1b). Additionally, the solutions from feasibility testing are used to reorient reversible reactions and further convert them into irreversible reactions, if applicable. The reorientation of reversible reactions minimizes the costly one-by-one step illustrated in Figure 1c. The structural analysis and dead-ends identification are performed each time a blocked reaction is identified during flux testing. The sign of LP7 solutions is used to reorient reversible reactions. Furthermore, as reversible exchange reactions are frequently involved in the one-by-one step, an additional step testing exchange reactions was added after the flux testing of irreversible reactions. Finally, in an optional step, the reversibility of the remaining reversible reactions is checked. A toy model in Figure 2 and the pseudocode in Box 1-4, further illustrate and detail the principle of FASTERCC.

**Figure 1:**
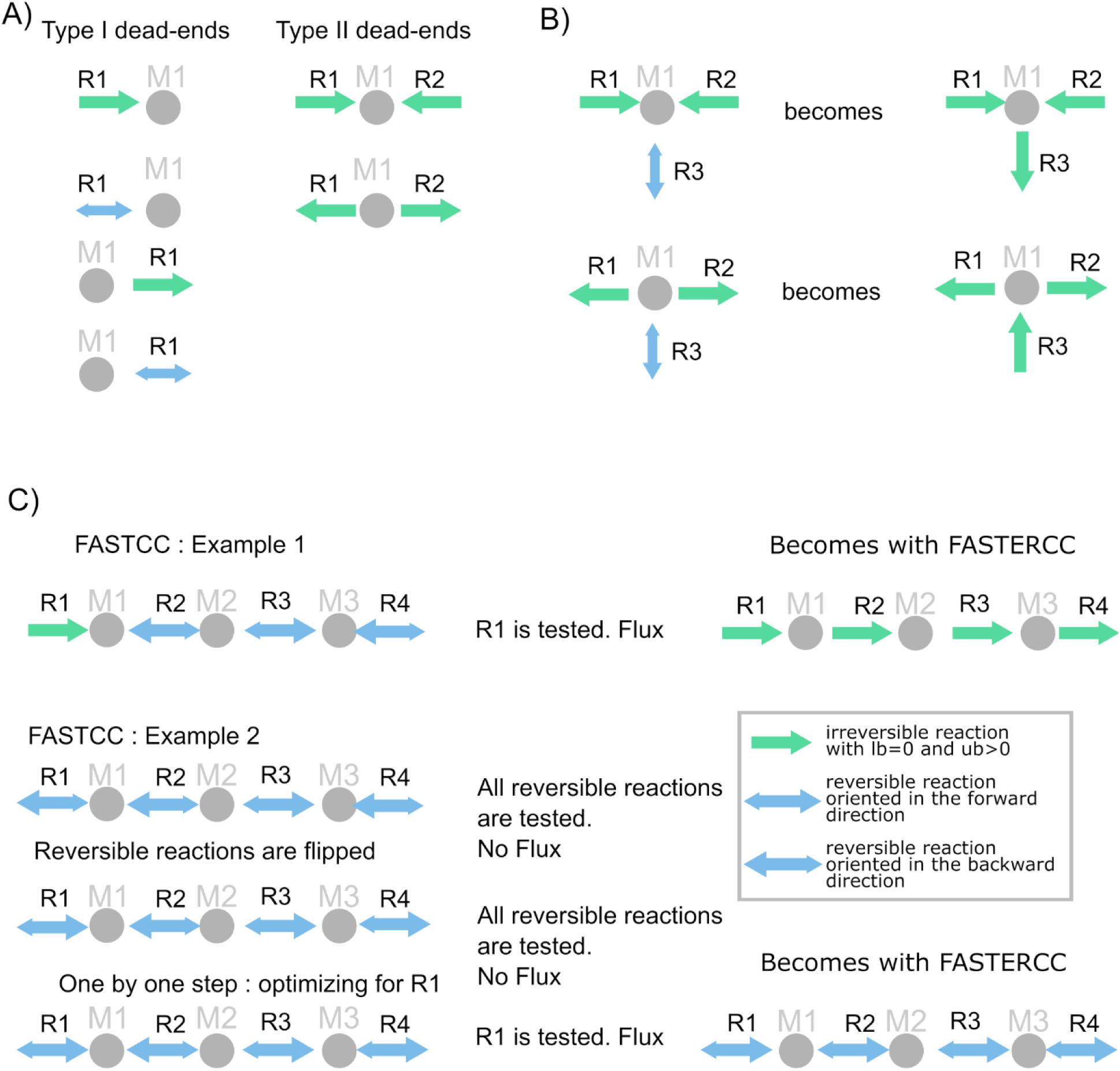
Speed-up strategies: Reversibility correction, dead-end detection, and reorientation of reversible reactions. Metabolites are illustrated as grey circles and labelled M1, M2, and M3. Irreversible and reversible reactions are depicted as green and blue arrows, respectively. The forward directions of reversible reactions (as defined in the stoichiometric matrix) are shown with larger arrowheads, while the smaller arrowheads depict the backward direction. A) FASTERCC identifies two types of dead-ends. Type I dead-ends are metabolites linked to only one reaction. Type II dead-ends are exclusively connected to only producing or only consuming irreversible reactions. Reactions linked to both dead-end types are removed during the structural analysis step (see Methods & Fig 2) B) If all reactions connected to a metabolite except one are irreversible producing or irreversible consuming reactions, then the reversible reaction is converted into an irreversible consuming reaction in the first example and into an irreversible producing reaction in the second example. C) FASTCC and FASTCORE evaluate the consistency of all irreversible reactions in the initial step (*e.g.*, R1 in example 1). To obtain a flux in R1, R2, R3, and R4 must carry flux. In example 2, where no irreversible reaction is present, all reversible reactions are tested simultaneously in one single LP. Due to their orientation, none can carry flux. Flipping all the reversible reactions does not resolve the situation. Only in the computationally expensive one-by-one step, when a single reaction is tested, a flux can be seen. Reorienting reversible reactions avoids this unnecessary flipping.

**Figure 2:**
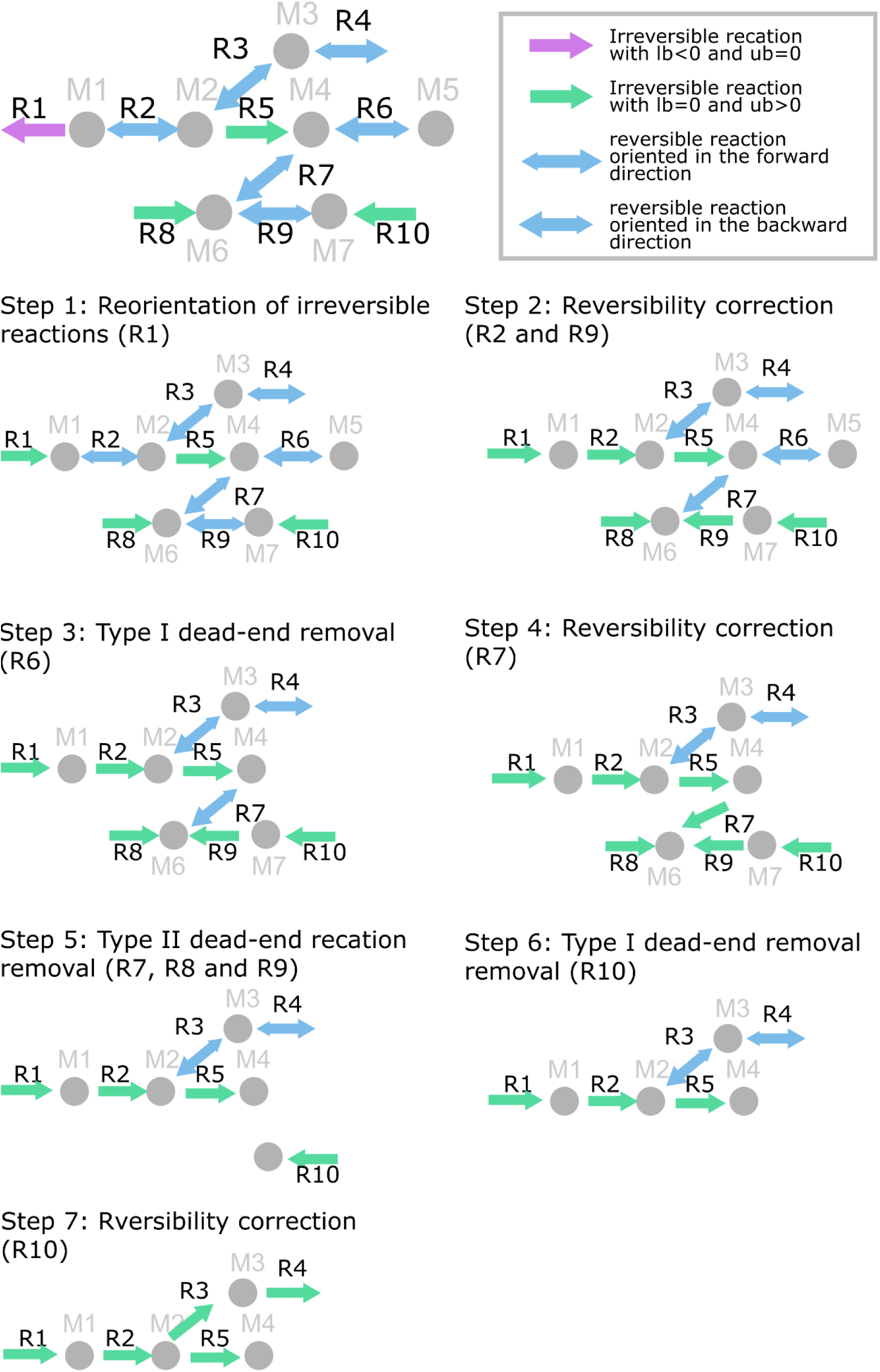
Overview of the structural analysis within FASTERCC: Within the structural analysis correcting reversibility of reaction and identifying and removing dead-ends is performed prior to any time-consuming feasibility test. The grey dots represent the metabolites; the arrows depict the reactions. Green arrows, blue, and pink arrows depict reversible, irreversible, and mis-oriented irreversible reactions. The sign in the stoichiometric matrix *S* is indicated by the larger head of the bidirectional arrows.

#### Box 1. The FASTERCC Algorithm for Flux Consistency Testing (short)

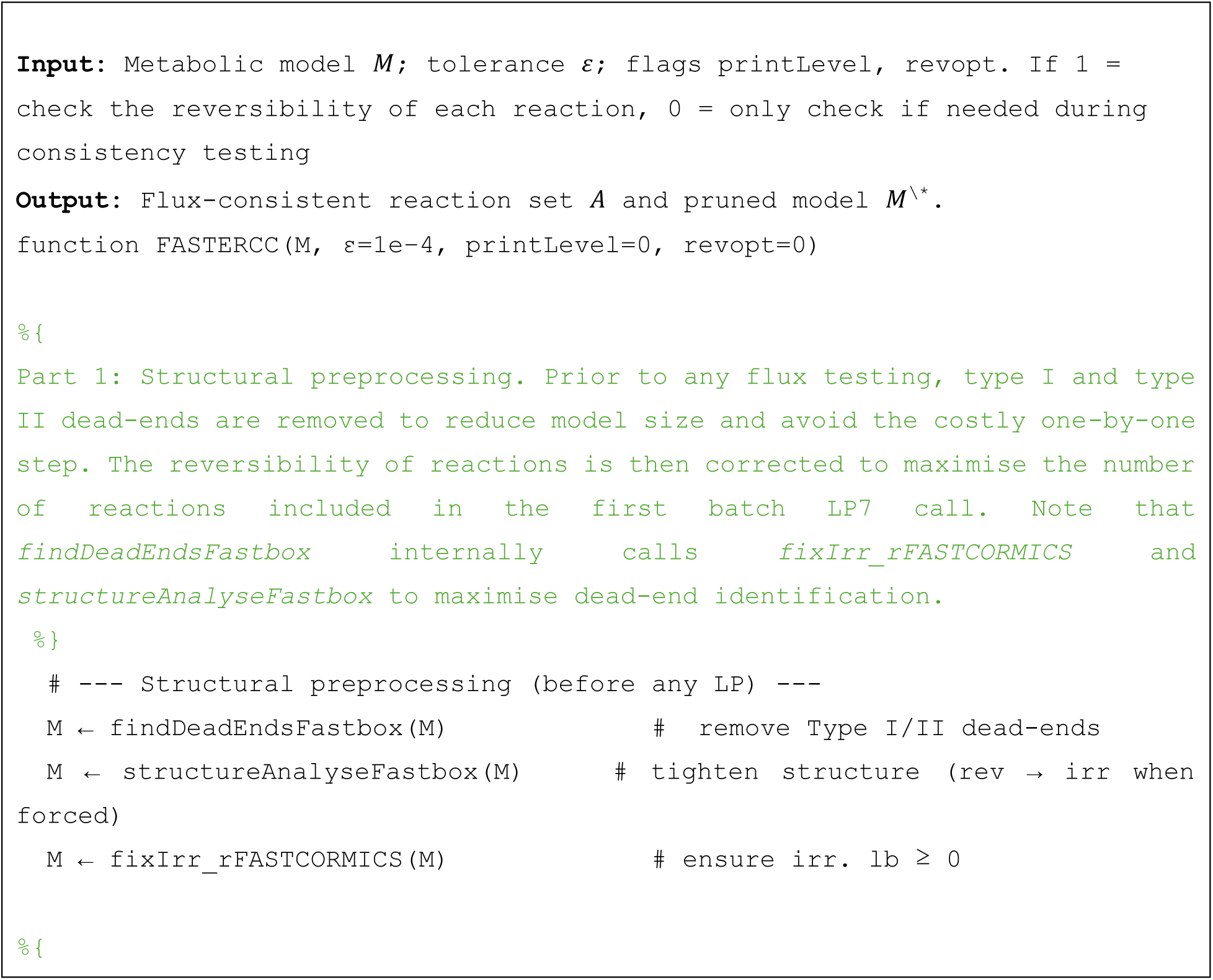
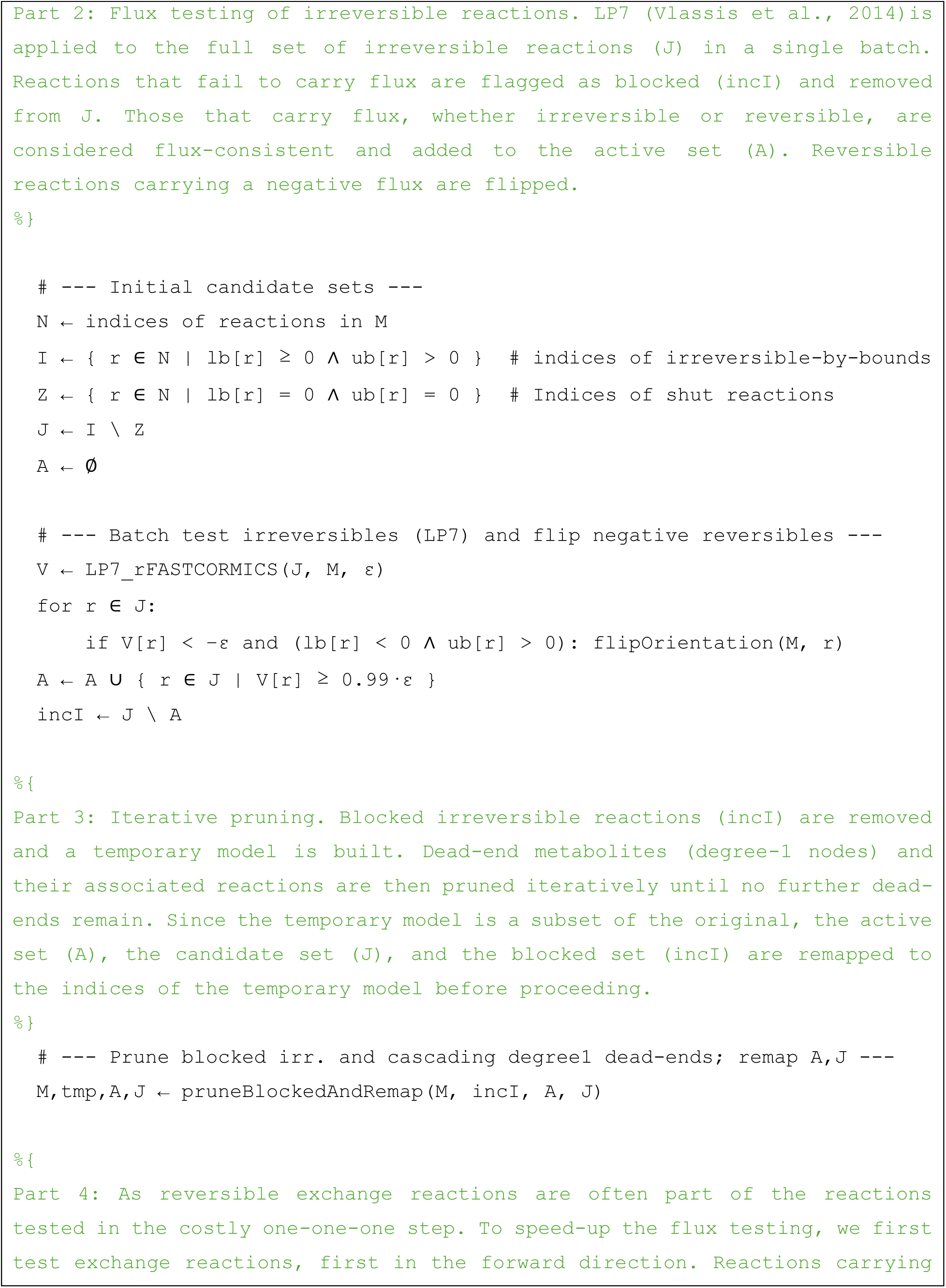
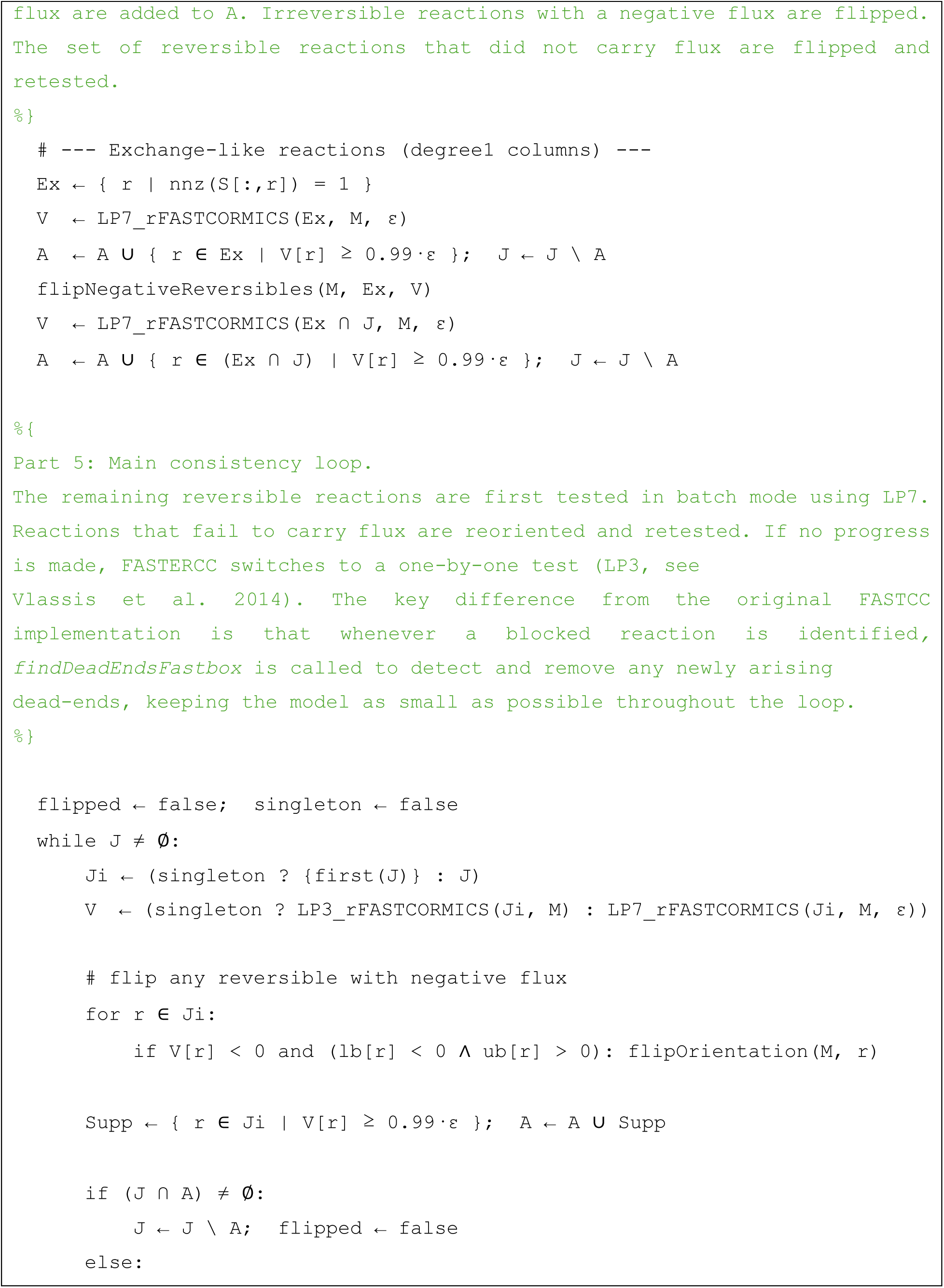
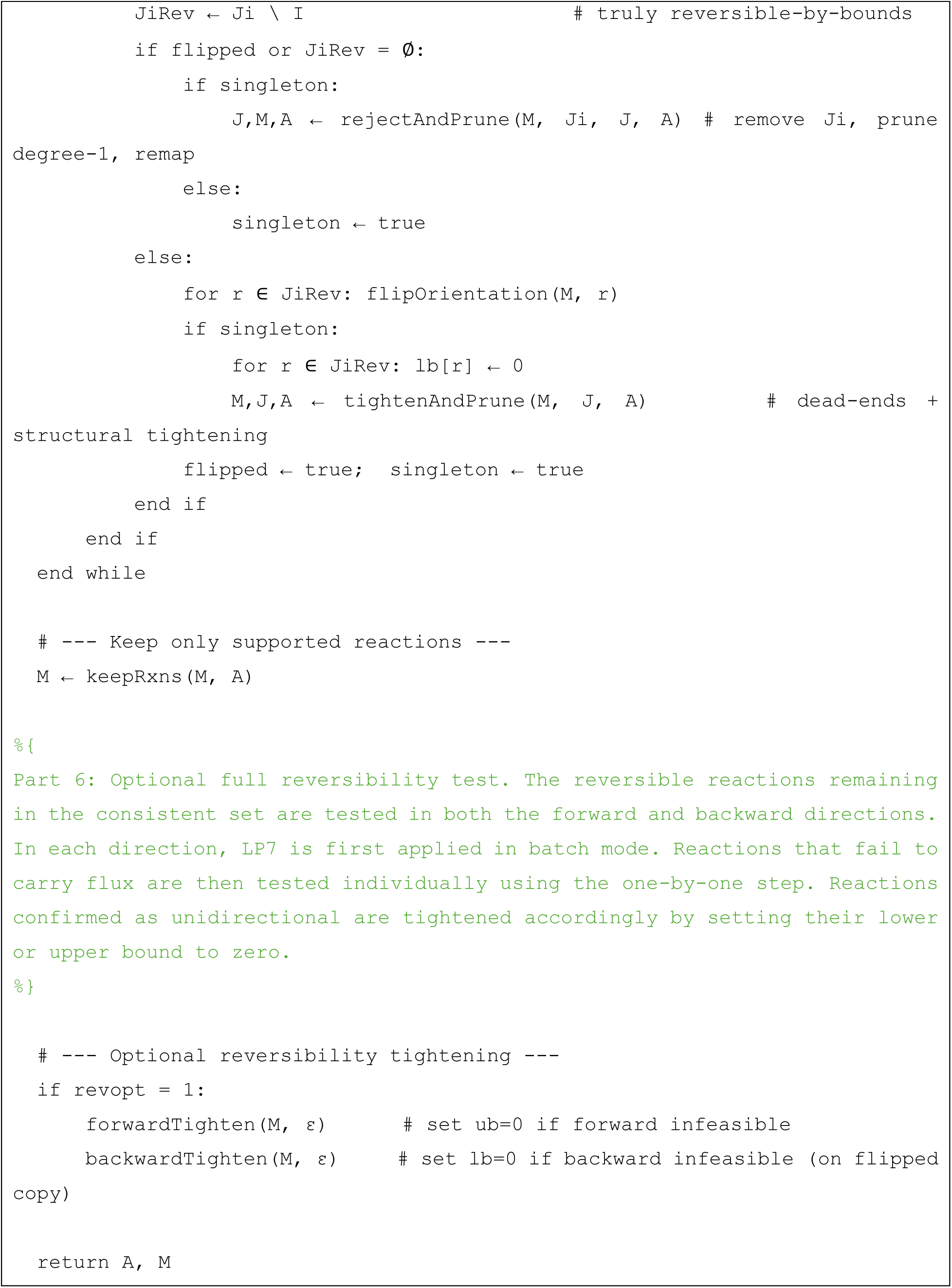
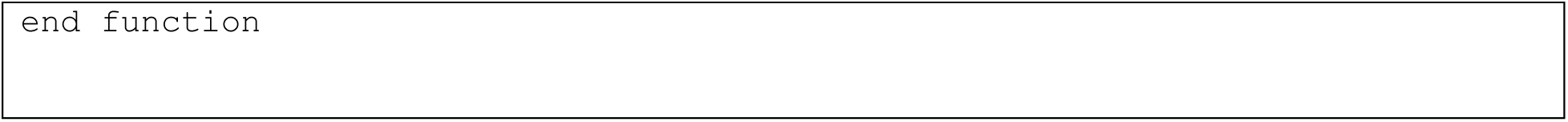

#### Box 2. The *fixIrr* function

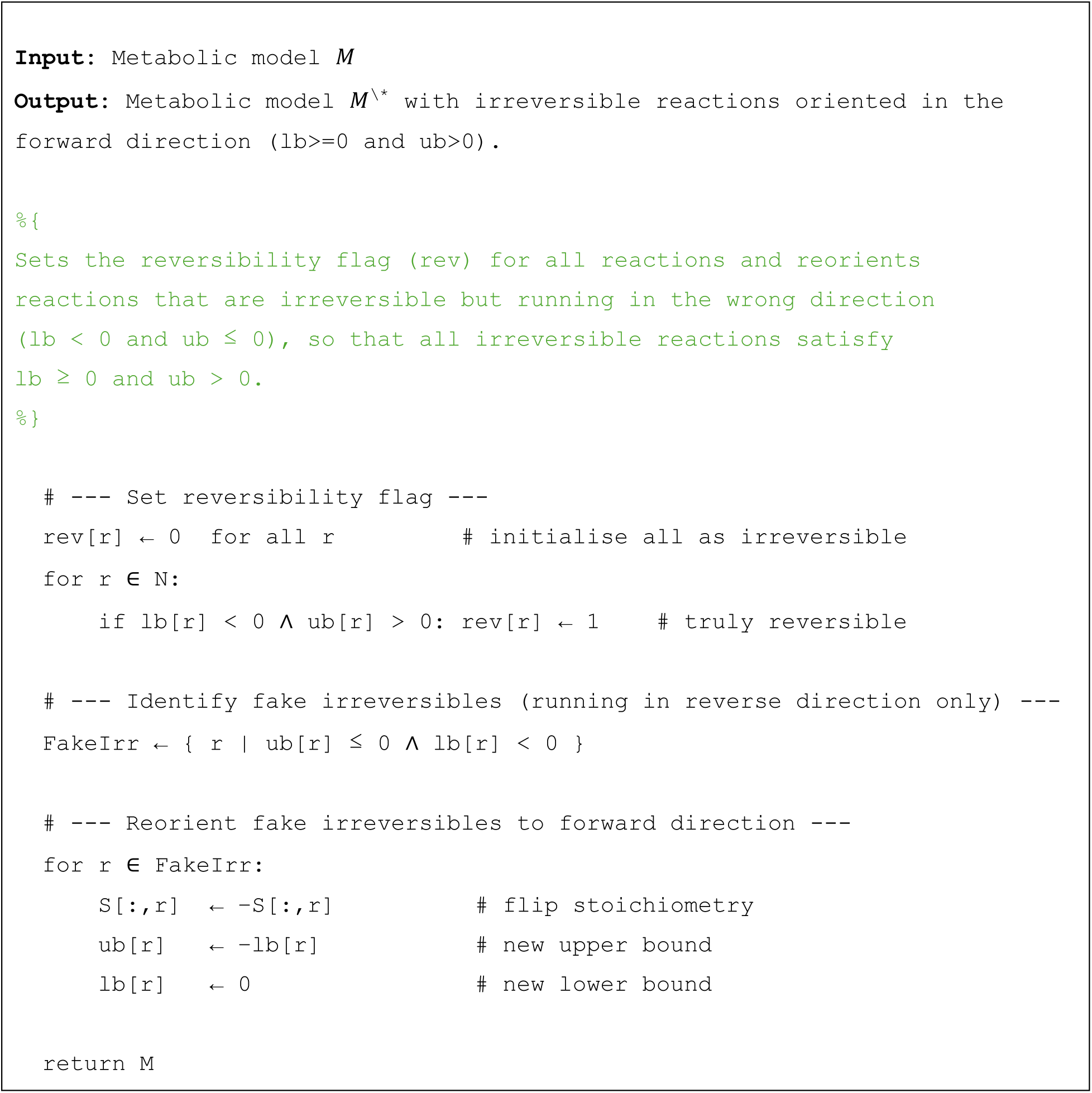

#### Box 3. The *structureAnalyseFastbox* function **Input:** Metabolic model *M*

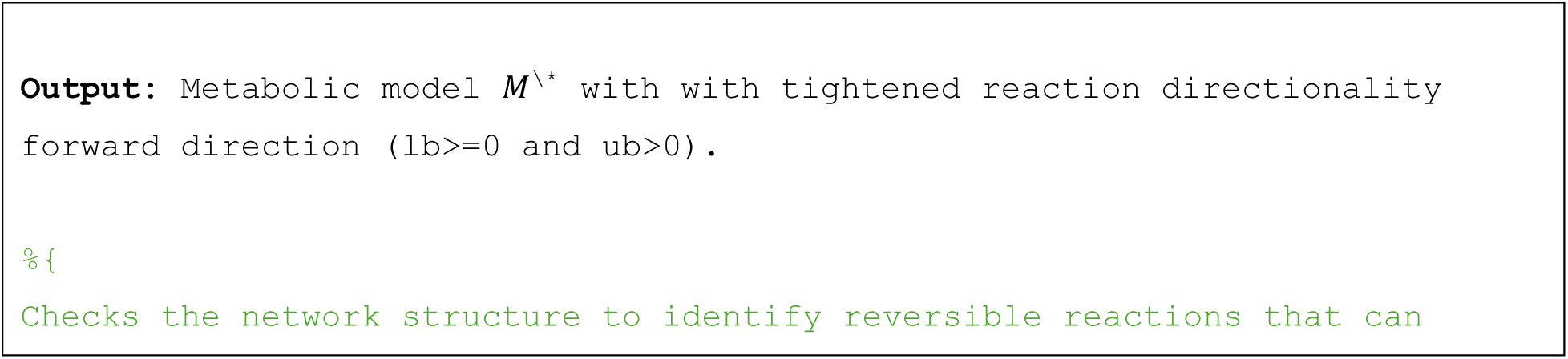
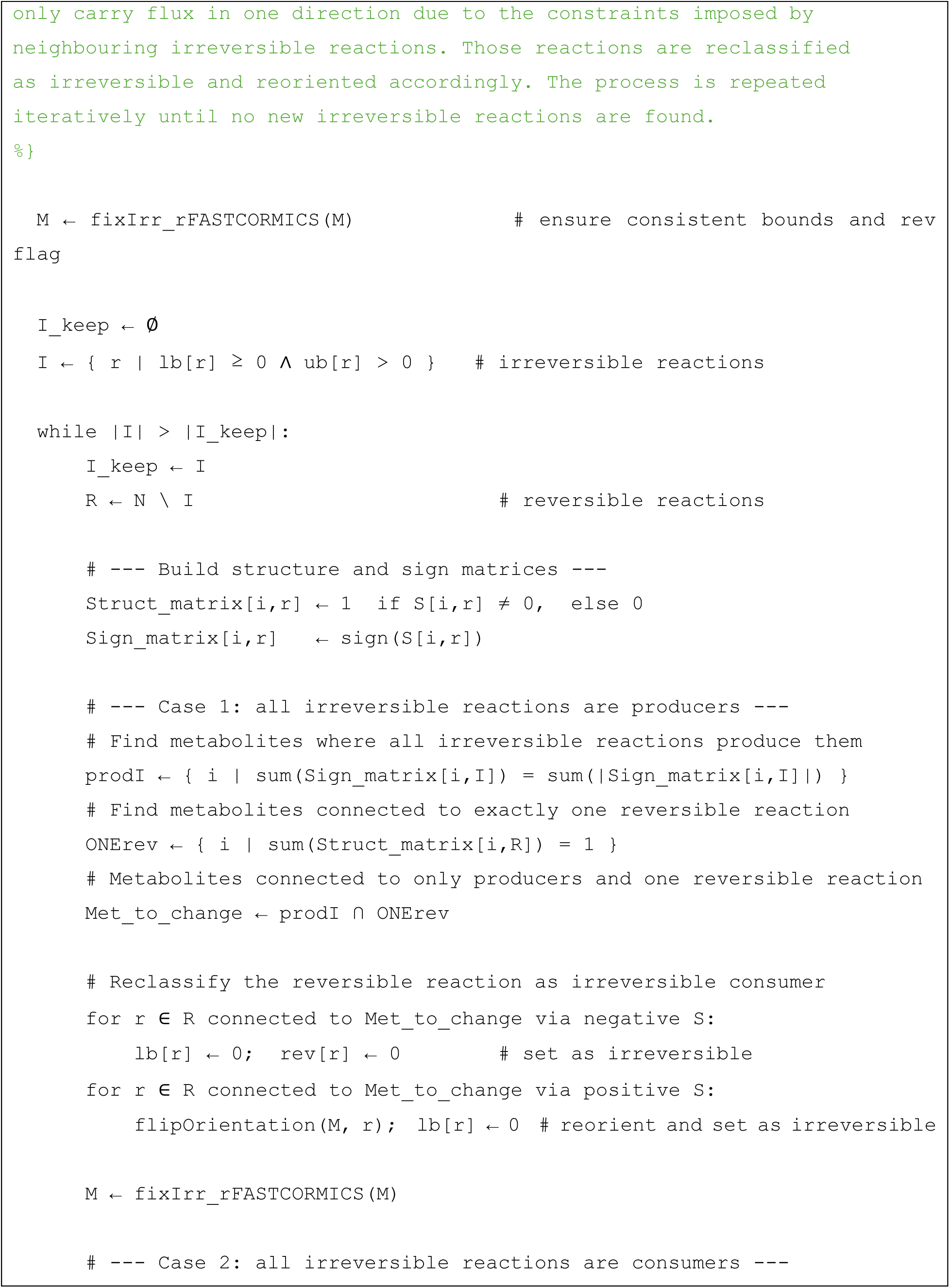
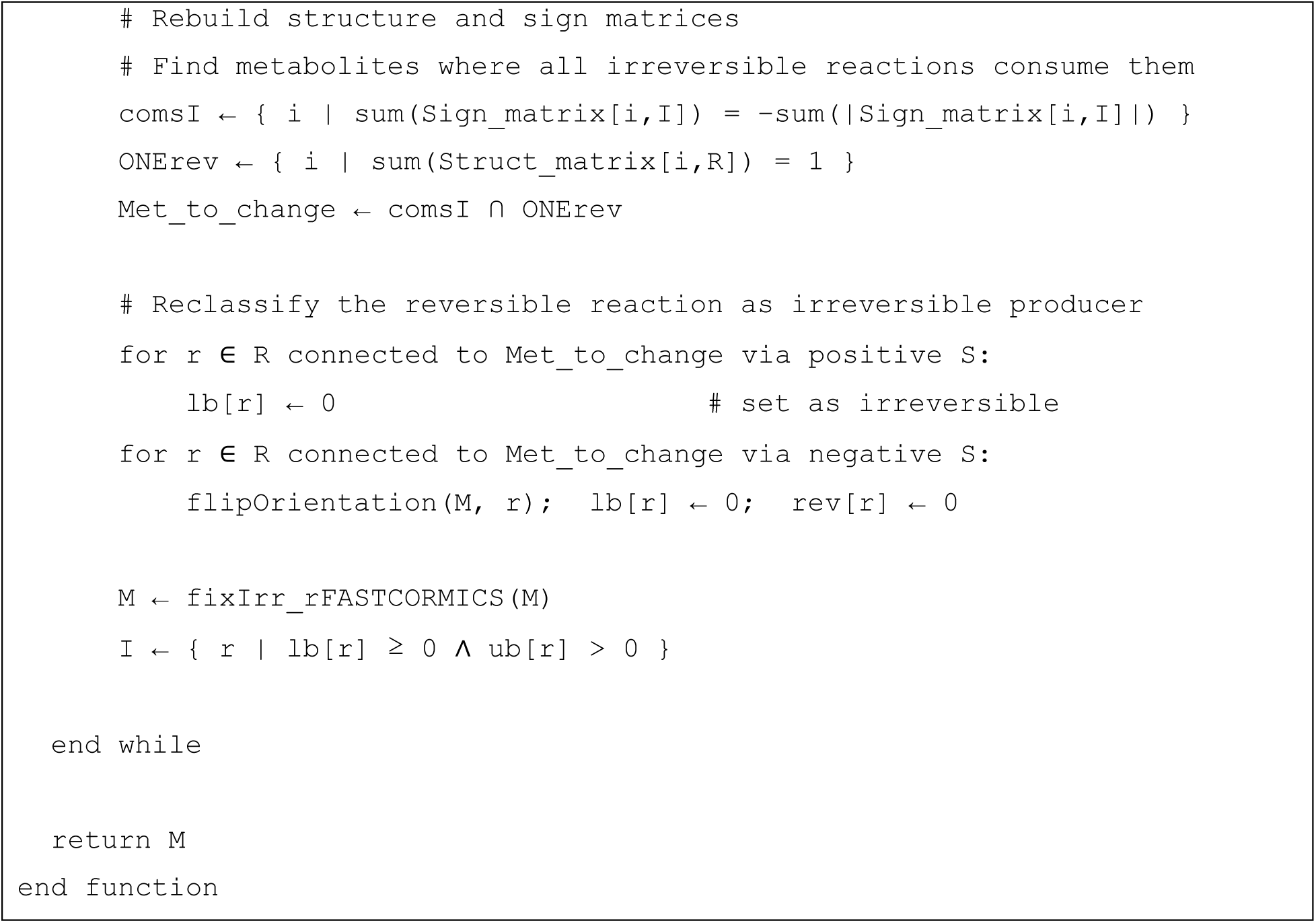

#### Box 4. The *findDeadEndsFastbox*

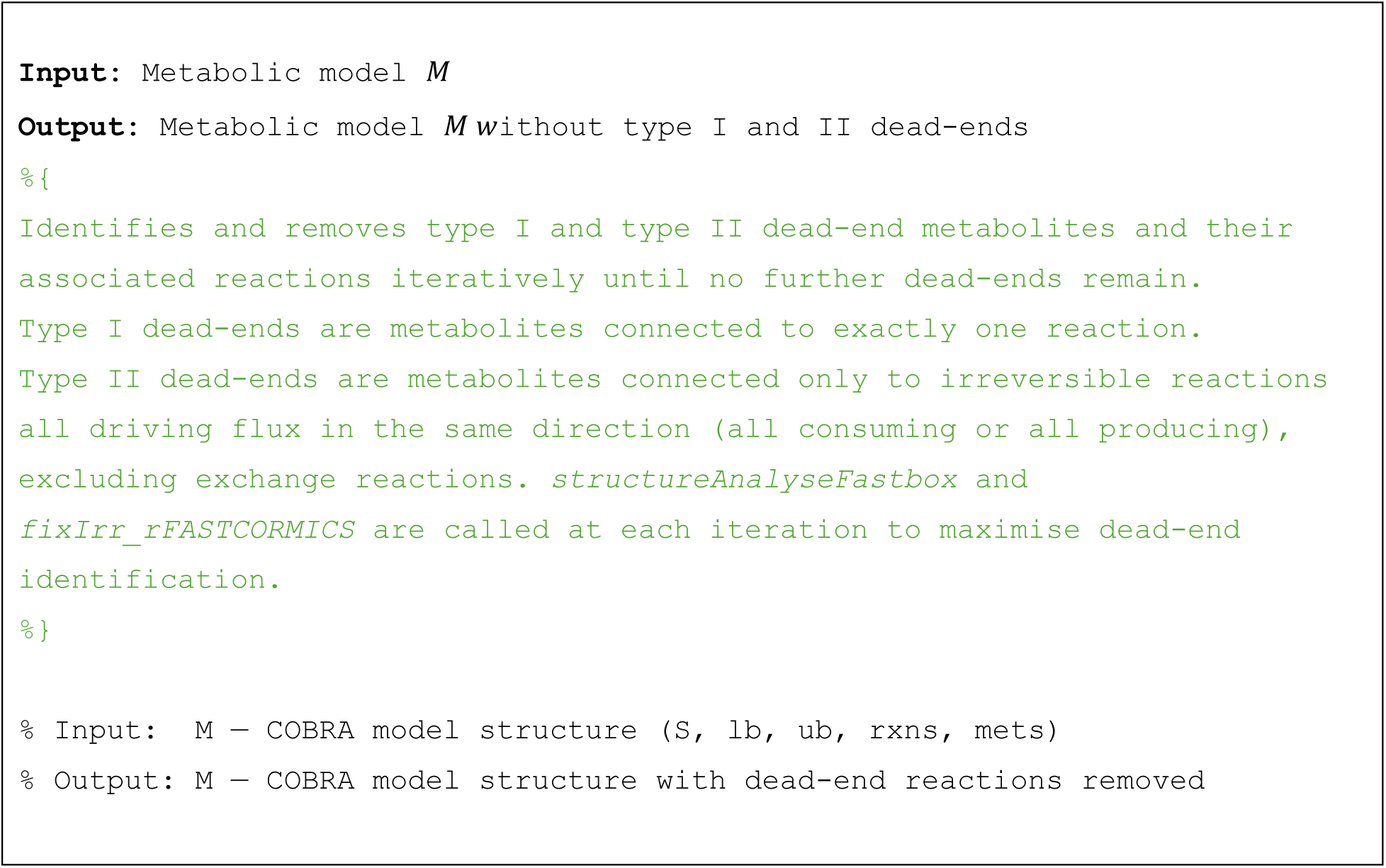
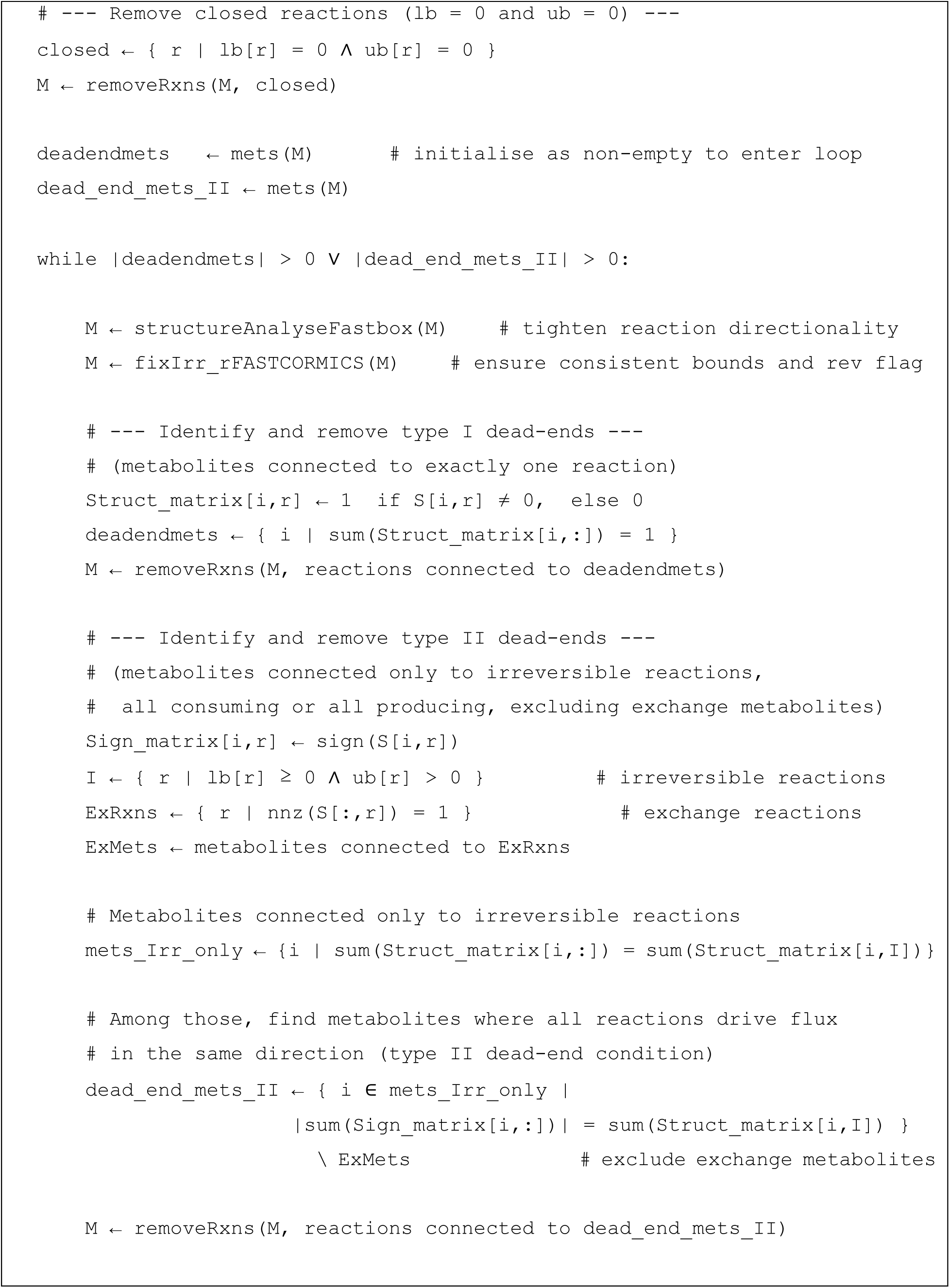
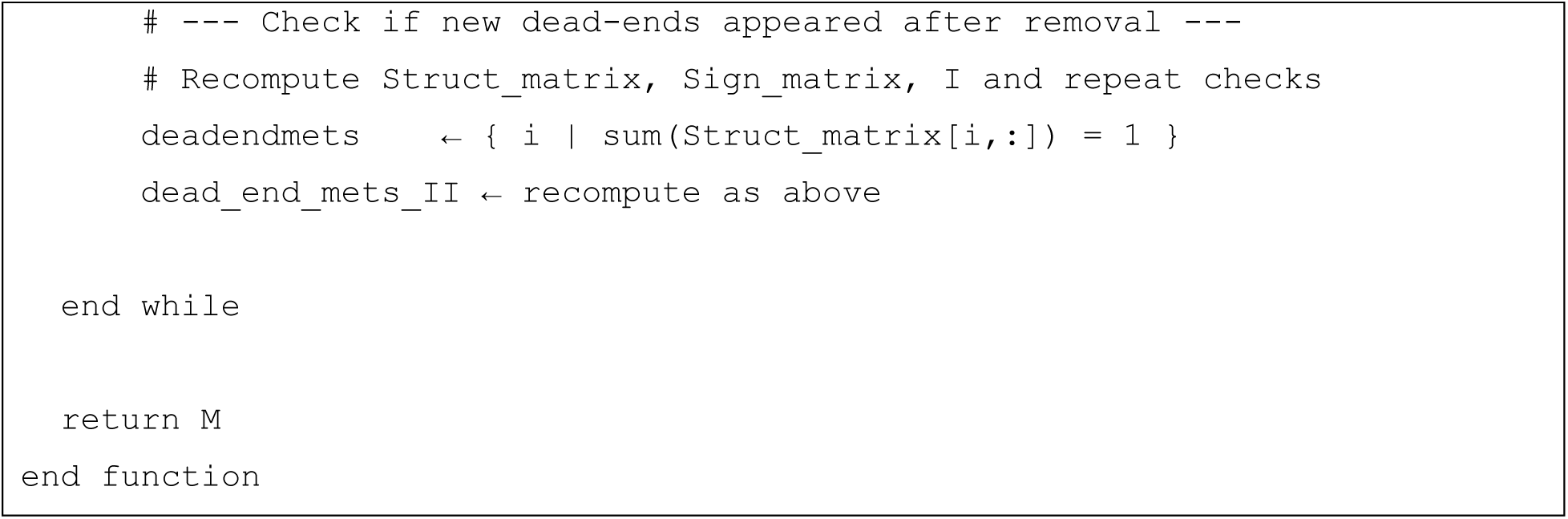

#### 1.1 Structural analysis

##### 1.1.1 Reorientation of irreversible reactions

In FASTERCC, first the sets of reversible and irreversible reactions are defined. Reversible reactions have a lower bound less than zero and an upper bound greater than zero. All remaining reactions are classified as irreversible. Irreversible reactions are reoriented (within the *fixIrr* function), if necessary, to have a non-negative lower bound. If a reaction *r* is reoriented (flipped), the signs of its coefficients in the stoichiometric matrix **S** are inverted, and its upper (*u*_*r*_) and lower (*l*_*r*_) bounds are swapped after multiplication by −1.

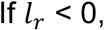

then the reaction is flipped:

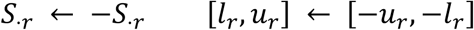

To ensure:

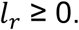

Reorientation is crucial as irreversible reactions with negative lower bounds can be wrongly classified as blocked by FASTCORE-based algorithms (Figure 2).

##### 1.1.2 Reversibility correction

A structural analysis is performed to minimize the number of total reversible reactions and hence necessary LPs. FASTERCC identifies reactions that appear reversible based on their bounds but are effectively irreversible within the network. For example, if a metabolite is connected to *r* reactions, of which *r − 1* are irreversible consuming reactions, then the remaining reversible reaction functions as an irreversible producing reaction and the bounds are updated accordingly. Similarly, if a metabolite is connected to r reactions where *r − 1* are irreversible producing reactions, the reversible reaction is effectively an irreversible consuming reaction (Figure 1). The process is repeated until no additional irreversible reaction is found (Figure 2).

##### 1.1.3 Dead-end detection

The reversibility correction is followed by dead-end detection. We identify type I dead-end metabolites as those connected to only one reaction (Figure 1). Once all reactions linked to type I dead-ends have been removed, the network is tested for type II dead-ends. Type II dead-ends are metabolites which are produced but not consumed or *vice versa*.

Let *Irr(m)* denote the set of irreversible reactions involving metabolite *m*, and let *S_mj_* be the stoichiometric coefficient of *m* in reaction *j*. A metabolite *m* is classified as a **Type II dead-end** if all irreversible reactions involving it have the same effect, either solely producing or solely consuming it:

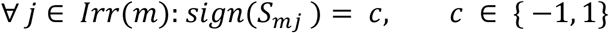

- c=+1: metabolite is exclusively *produced*
- c=-1: metabolite is exclusively *consumed*

Equivalently, this condition can be expressed as

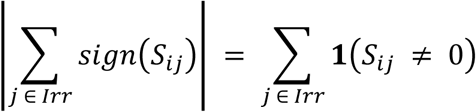

where S is the stoichiometric matrix and *S*_*ij*_ denotes the stoichiometric metric coefficients of metabolite i in reaction j. The *sign*(*S*_*ij*_) returns 1 if the metabolite is produced and -1 if consumed and 0 if the metabolite does not participate in the reaction. The **1**(*S*_*ij*_ ≠ 0) represent the structural matrix and hence indicates if a metabolite i participates or not in a reaction. The equality finds the instances where all the reactions r connected to i are exclusively positive or negative (type II dead-end).

Reactions linked to type I and II dead-ends are removed iteratively. As the identification of type II dead-ends requires that all connected reactions to be irreversible, and as irreversible reactions are central for the reversibility corrections, the structural analysis is rerun each time a new irreversible reaction is identified during the feasibility testing.

#### 1.2 Reorientation of reversible reactions and directionality correction

Reactions that during feasibility testing had a negative flux are reoriented. Reactions that failed to carry flux in either the forward or the backward direction but carry a flux in the other direction are converted to the respective irreversible reactions.

### 2. Benchmarking

FASTERCC and FASTCC were tested under the same settings: a standard laptop with 4 CPU cores and 16 GB RAM and on an HPC (Varrette et al., 2014) with 52224 GB RAM for larger networks and number of runs. Both algorithms were run using CPLEX and with an epsilon of 1e-4. All experiments were performed in MATLAB R2019b to avoid compatibility issues between MATLAB and CPLEX. To ensure a fair comparison, no parallelization was used.

To systematically compare FASTCC and FASTERCC across a wide range of network sizes and numbers of blocked reactions, we created 20 expanded models composed of 1–10 clusters connected to a union compartment to the environment using the function *build_expanded_input_mode*l of scFASTCORMICS (Pacheco et al., 2022). Each cluster consists of a copy of 10,600 reactions from Recon3D (Brunk et al., 2018). Then, to simulate blocked reactions, 10, 50, 100, 200, or 500 reactions were randomly removed from these input models, and FASTCC and FASTERCC were then applied to identify the set of consistent reactions. The size of the consistent models obtained by both algorithms and the running time were recorded. We also tested FASTCC (Vlassis et al., 2014) and FASTERCC against *SwiftCC* (Tefagh & Boyd, 2020), but *SwiftCC* overestimates the number of flux-consistent reactions declaring some reactions that carry zero fluxes according to Flux Variability Analysis (FVA) as consistent and hence was not included here.

Expanded models were reconstructed using the same protocol as in the previous experiment. Core sets were randomly generated and FASTCORE was then run to reconstruct context-specific models with and without FASTERCC as preprocessing step. FASTERCC has an option to correct the reversibility of all the reactions in a model. We validated the results using the *fluxVariability* function of the COBRA Toolbox (Heirendt et al., 2019). Reactions found by this function with a lower range below -epsilon (*-1e-4*) and an upper range above epsilon (*1e-4*) were considered reversible, while the remaining reactions were categorised as irreversible.

## Results

To assess the scalability of FASTERCC and FASTCC, we tested the consistency of 1140 models, ranging from 21,200 to 183,920 reactions, with varying numbers of blocked reactions (0-500). Across all instances, FASTCC and FASTERCC identified identical sets of flux consistent reactions (data not shown). However, the mean speedup of FASTERCC was between 2.3 and 31.5 times faster, especially for larger networks with a high number of blocked reactions. For example, when 500 blocked reactions were present, the speedup ranged from 17.5-fold to 31.5-fold across model sizes, whereas for consistent networks (0 blocked reactions) the speedup ranged from 2.3-fold to 5.2-fold. While FASTCC running times grew substantially with model size and number of blocked reactions, the running times of FASTERCC remained below 1,800 seconds (≈30 minutes) even for larger networks (see Figure 3 and Table 1). On consistent networks, FASTERCC was 2.3 to 5.2 times faster than FASTCC (Vlassis et al., 2014) (see Figure 3 and Table 1). The detailed mean running times of FASTCC and FASTERCC, as well as the speedups for the various network sizes and numbers of removed reactions, can be found in Supplementary Table 1.

**Figure 3.**
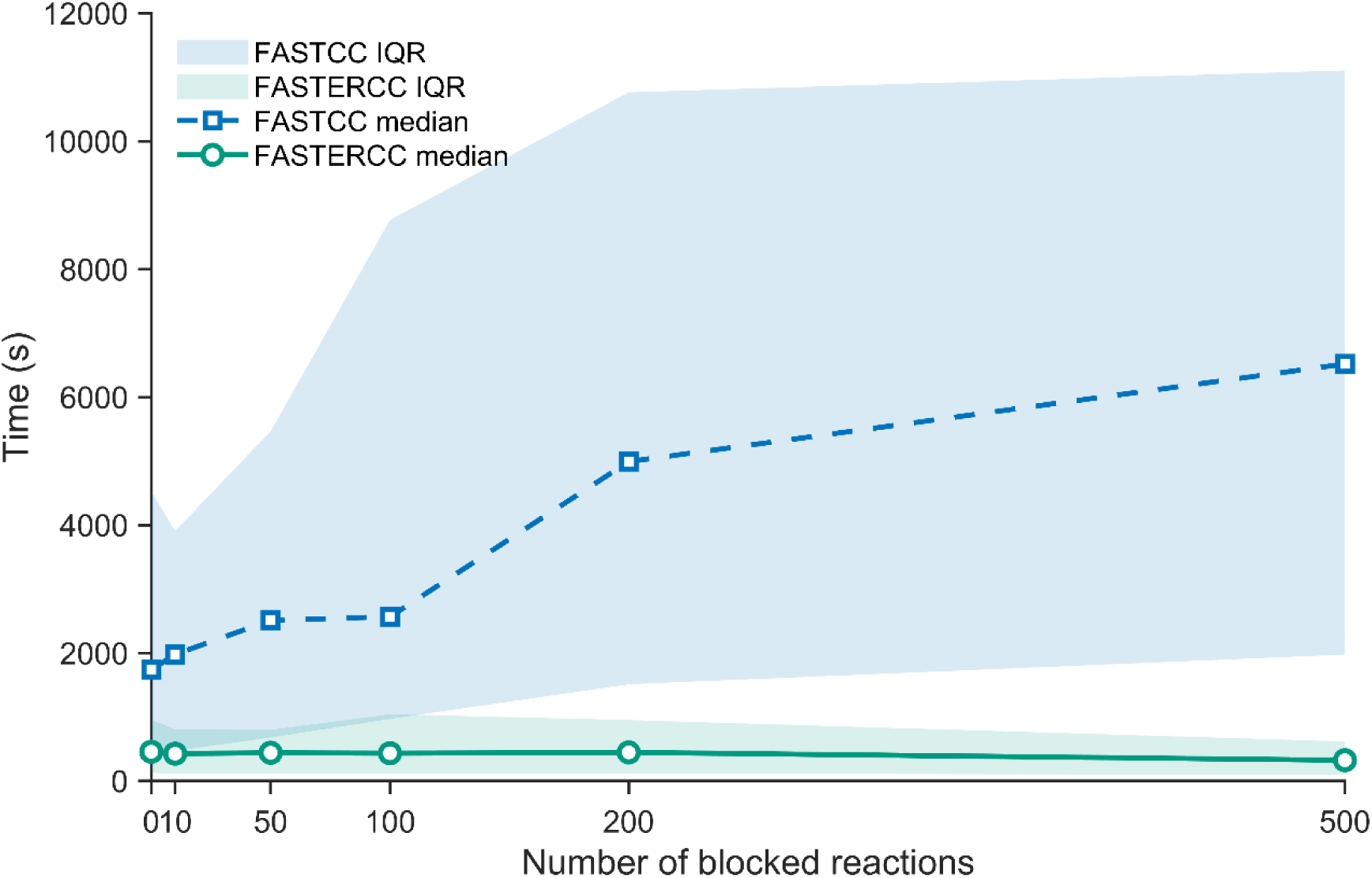
Runtime comparison of FASTCC and FASTERCC across networks with increasing numbers of blocked reactions. Median runtimes of FASTCC (blue dashed line) and FASTERCC (green line) are shown for benchmark networks containing 0–500 blocked reactions. Shaded areas indicate the interquartile range (IQR) across cluster configurations.

**Table 1:** Comparison of the running time of FASTCC and FASTERCC across model sizes and percentages of blocked reactions.

| *Input Model Size* | *Number of blocked reactions* | *FASTCC Time (in sec)* | *FASTERCC Time (in sec)* | *Speedup* | *Same result* |
| --- | --- | --- | --- | --- | --- |
| 21200 | 500 | 655.2 | 20.8 | **31.5** | Yes |
| 30240 | 500 | 1033.3 | 33.9 | **30.5** | Yes |
| 39280 | 500 | 1723.8 | 68.9 | **25.0** | Yes |
| 57360 | 500 | 2940.6 | 125.3 | **23.5** | Yes |
| 84480 | 500 | 10790.0 | 499.6 | **21.6** | Yes |
| 75440 | 500 | 4881.6 | 237.7 | **20.5** | Yes |
| 93520 | 500 | 11107.7 | 559.2 | **19.9** | Yes |

To assess if the reversibility correction and the reorientation of reversible reactions could improve the performance of FASTCORE, we reconstructed 225 consistent networks from 10,600 to 93,520 reactions and various number of core reactions and run FASTCORE (Vlassis et al., 2014) with and without preprocessing with FASTERCC (see Figure 4 and Table 2). A speed-up of 1.8-fold could especially be observed for larger networks. Overall, FASTCORE runtime without preprocessing ranged from 4.4 s to 586.5 s, whereas runtime with FASTERCC preprocessing ranged from 2.7 s to 475.1 s (Figure 4 and Table 2). The preprocessing reduced the running time of FASTCORE in 183 out of 250 runs by at least 10% and with a top speed up of 3-fold. The detailed mean running times of FASTCORE with and without FASTERCC preprocessing, as well as the speedups for the various network sizes and numbers of core reactions, can be found in Supplementary Table 2.

**Figure 4.**
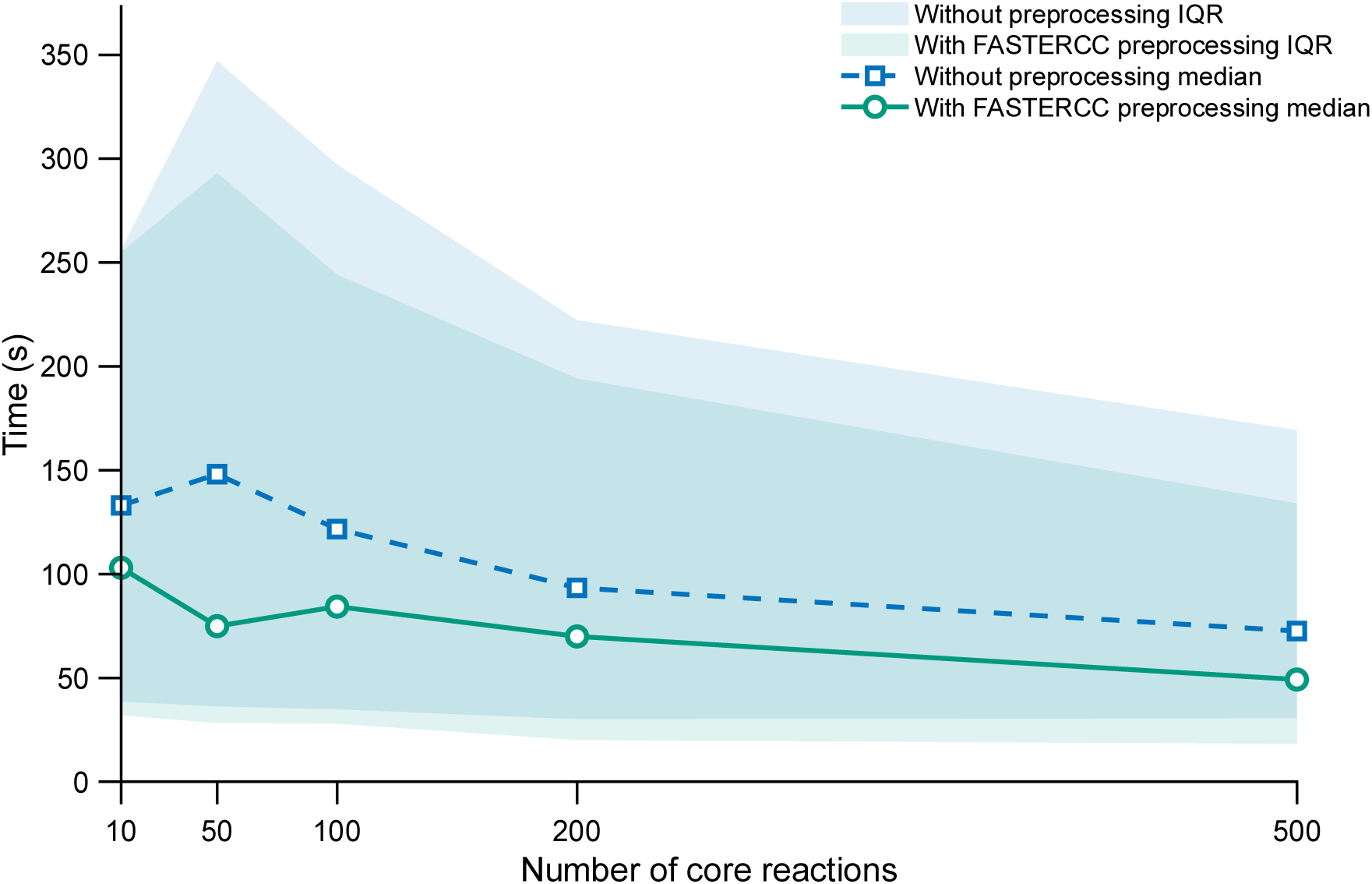
Runtime comparison of FASTCORE with and without FASTERCC preprocessing across increasing core reaction set sizes. Median runtimes of FASTCORE without preprocessing (blue dashed line) and FASTCORE with FASTERCC preprocessing (green line) are shown for benchmark networks with core reaction sets ranging from 10 to 500 reactions. Shaded areas indicate the interquartile range (IQR) across cluster configurations.

**Table 2:** FASTCORE running time improvement due to the preprocessing with FASTERCC.

| *Input Model Size* | *Number of Core reactions* | *Time (No preprocessing)* | *Time with FASTERCC preprocessing* | *Speedup* |
| --- | --- | --- | --- | --- |
| 48320 | 200 | 118.2 | 65.4 | 1.8 |
| 84480 | 500 | 481.5 | 276.0 | 1.7 |
| 84480 | 200 | 480.6 | 299.3 | 1.6 |
| 21200 | 100 | 17.9 | 10.8 | 1.7 |
| 57360 | 10 | 121.6 | 80.1 | 1.5 |

FASTERCC can optionally check assess reversibility of each reaction in the network and convert reversible reactions constrained by the network topology to carry flux exclusively in the forward reaction or backward direction into irreversible reactions. The accuracy of the reversibility correction was confirmed with flux variability analysis (FVA) on Recon3D (Brunk et al., 2018), ihuman1 (Robinson et al., 2020), iMM1415 (Sigurdsson et al., 2010) and iMM1865 (Khodaee et al., 2020), see Table 3.

**Table 3:** Number of reversible reactions based on bounds and network topology.

| ***Model*** | ***Number of reversible reactions (bounds)*** | ***Number of reversible reactions (FASTERCC-network)*** | ***Number of reversible reactions (FVA-network)*** |
| --- | --- | --- | --- |
| *Recon3D* | 5234 | 3520 | 3520 |
| *lhuman (Human-GEM v1.13.0)* | 5975 | 3594 | 3594 |
| *iMM1415* | 1185 | 363 | 363 |
| *iMM1865* | 5244 | 3531 | 3531 |

## Discussion

Low computational times are not just a matter of convenience. They enable a whole set of *in silico* experiments such as parameter optimizations, large scale sensitivity analysis, and cross-validation workflows requiring multiple model rebuilds. These experiments were infeasible for models exceeding 100,000 reactions without access to HPC (Varrette et al., 2014). Even with HPC, running longer experiments with varying epsilon values, expression thresholds, and core sets can be challenging with wall-time limits, maintenance, and resource contention.

In this paper, we extend FASTCC (Vlassis et al., 2014) into FASTERCC, a scalable algorithm that not only accelerates flux consistency testing but, through the correction of reversibility and orientation of reactions also speeds up FASTCORE (Vlassis et al., 2014), a central tool in metabolic modelling. While FASTCC, FASTERCC and FVA consistently give the same reactions. FASTCORE can build slightly different models due to different optima. The preprocessing of FasterCC is expected to benefit other FASTCORE-based workflows, toolboxes, and pipelines FASTCORMICS (Pacheco et al., 2015b), rFASTCORMICS (Pacheco et al., 2019), scFASTCORMICS (Pacheco et al., 2022), Cobra toolbox (Heirendt et al., 2019), FastGapFill (Thiele et al., 2014), PSAMM (Steffensen et al., 2016), TROPPO (Cunha et al., 2023), COMO (Bessell et al., 2023), TRFBA-CORE (Jamialahmadi et al., 2019), ThermOptiCS (Kumar S & Bhatt, 2025) Finally, correcting reversibility using FASTERCC has the potential to reduce computational times of workflows such as flux variability analysis, which tests reversible reactions in the forward and the backward direction. It could also benefit workflows that split reversible reactions in two irreversible ones by reducing reaction counts and the limiting loop formation between the forward and backward reactions causing the appearance of blocked reactions. FASTERCC thus provides a foundation for scalable metabolic modelling in the era of large and complex biological datasets.

## Acknowledgement

The computational experiments presented in this paper were partially carried out using the HPC facilities of the University of Luxembourg, https://hpc.uni.lu (Varrette et al., 2014).

## Code availability

The MATLAB implementation of FASTERCC is available at: https://github.com/sysbiolux/rFASTCORMICS

## Supplementary File

**Table 1:** Comparison of the running time of FASTCC and FASTERCC across model sizes and percentages of blocked reactions. Input model sizes ranged from **21,200 to 183,920 reactions**. The number of blocked reactions ranged from **0 to 500**. In all reported cases, FASTCC and FASTERCC returned the same flux-consistent reaction set.

| Input Model Size | Number of blocked reactions | FASTCC time mean | FASTCC Time SD | FasterCC Time mean | FasterCC Time SD | Speedup from means | Same Result |
| --- | --- | --- | --- | --- | --- | --- | --- |
| 21200 | 0 | 82.4 | 33.2 | 18.6 | 14.6 | 4.4 | 1 |
| 21200 | 10 | 55.7 | 9.8 | 10.5 | 1.2 | 5.3 | 1 |
| 21200 | 50 | 100.6 | 25.9 | 10.6 | 1.1 | 9.5 | 1 |
| 21200 | 100 | 196.4 | 47.2 | 13.3 | 1.8 | 14.8 | 1 |
| 21200 | 200 | 307.4 | 46.0 | 13.7 | 2.1 | 22.4 | 1 |
| 21200 | 500 | 655.2 | 59.6 | 20.8 | 2.0 | 31.5 | 1 |
| 30240 | 0 | 161.5 | 44.6 | 43.9 | 41.2 | 3.7 | 1 |
| 30240 | 10 | 115.4 | 8.5 | 24.1 | 1.8 | 4.8 | 1 |
| 30240 | 50 | 190.6 | 44.4 | 27.3 | 3.7 | 7.0 | 1 |
| 30240 | 100 | 365.0 | 91.2 | 27.2 | 2.9 | 13.4 | 1 |
| 30240 | 200 | 507.2 | 83.1 | 30.4 | 2.5 | 16.7 | 1 |
| 30240 | 500 | 1033.3 | 184.0 | 33.9 | 4.0 | 30.5 | 1 |
| 39280 | 0 | 234.5 | 96.3 | 60.8 | 38.9 | 3.9 | 1 |
| 39280 | 10 | 271.0 | 122.1 | 57.0 | 5.9 | 4.8 | 1 |
| 39280 | 50 | 359.7 | 57.7 | 57.9 | 4.9 | 6.2 | 1 |
| 39280 | 100 | 571.0 | 109.3 | 60.5 | 6.3 | 9.4 | 1 |
| 39280 | 200 | 860.6 | 196.3 | 59.8 | 10.0 | 14.4 | 1 |
| 39280 | 500 | 1723.8 | 347.0 | 68.9 | 14.2 | 25.0 | 1 |
| 48320 | 0 | 284.1 | 125.6 | 102.7 | 52.6 | 2.8 | 1 |
| 48320 | 10 | 321.2 | 107.7 | 75.3 | 5.6 | 4.3 | 1 |
| 48320 | 50 | 424.8 | 58.5 | 81.4 | 12.1 | 5.2 | 1 |
| 48320 | 100 | 639.6 | 174.5 | 78.6 | 7.9 | 8.1 | 1 |
| 48320 | 200 | 1012.9 | 148.0 | 84.4 | 5.2 | 12.0 | 1 |
| 48320 | 500 | 1976.8 | 227.4 | 92.9 | 14.9 | 21.3 | 1 |
| 57360 | 0 | 458.0 | 162.8 | 115.5 | 34.9 | 4.0 | 1 |
| 57360 | 10 | 425.3 | 56.9 | 103.6 | 6.6 | 4.1 | 1 |
| 57360 | 50 | 629.3 | 110.8 | 100.5 | 14.1 | 6.3 | 1 |
| 57360 | 100 | 948.4 | 137.3 | 106.2 | 10.4 | 8.9 | 1 |
| 57360 | 200 | 1400.6 | 321.4 | 108.4 | 7.1 | 12.9 | 1 |
| 57360 | 500 | 2940.6 | 493.1 | 125.3 | 21.1 | 23.5 | 1 |
| 66400 | 0 | 579.4 | 184.8 | 160.9 | 38.0 | 3.6 | 1 |
| 66400 | 10 | 614.2 | 109.8 | 140.5 | 14.7 | 4.4 | 1 |
| 66400 | 50 | 834.8 | 253.4 | 150.1 | 12.6 | 5.6 | 1 |
| 66400 | 100 | 1041.1 | 210.2 | 141.3 | 21.4 | 7.4 | 1 |
| 66400 | 200 | 1849.5 | 229.7 | 152.6 | 21.5 | 12.1 | 1 |
| 66400 | 500 | 3723.3 | 583.3 | 165.5 | 10.5 | 22.5 | 1 |
| 75440 | 0 | 742.3 | 143.9 | 327.3 | 414.2 | 2.3 | 1 |
| 75440 | 10 | 773.5 | 36.2 | 177.4 | 13.1 | 4.4 | 1 |
| 75440 | 50 | 1116.1 | 111.6 | 177.1 | 17.5 | 6.3 | 1 |
| 75440 | 100 | 1720.0 | 508.0 | 212.0 | 22.1 | 8.1 | 1 |
| 75440 | 200 | 2423.9 | 418.3 | 220.0 | 48.8 | 11.0 | 1 |
| 75440 | 500 | 4881.6 | 590.6 | 237.7 | 21.6 | 20.5 | 1 |
| 84480 | 0 | 1151.0 | 462.2 | 255.3 | 62.5 | 4.5 | 1 |
| 84480 | 10 | 961.6 | 36.3 | 242.8 | 12.7 | 4.0 | 1 |
| 84480 | 50 | 1279.9 | 132.2 | 238.6 | 32.0 | 5.4 | 1 |
| 84480 | 100 | 1863.3 | 369.0 | 238.2 | 25.7 | 7.8 | 1 |
| 84480 | 200 | 5082.3 | 669.0 | 447.8 | 94.6 | 11.3 | 1 |
| 84480 | 500 | 10790.0 | 1078.4 | 499.6 | 57.1 | 21.6 | 1 |
| 93520 | 0 | 1308.6 | 184.3 | 457.8 | 406.3 | 2.9 | 1 |
| 93520 | 10 | 1321.0 | 91.0 | 302.0 | 30.5 | 4.4 | 1 |
| 93520 | 50 | 1807.1 | 254.4 | 279.9 | 31.8 | 6.5 | 1 |
| 93520 | 100 | 2294.1 | 335.6 | 293.4 | 53.3 | 7.8 | 1 |
| 93520 | 200 | 3680.3 | 513.4 | 310.8 | 34.9 | 11.8 | 1 |
| 93520 | 500 | 11107.7 | 1029.6 | 559.2 | 97.8 | 19.9 | 1 |
| 102560 | 0 | 1740.3 | 422.5 | 388.3 | 83.8 | 4.5 | 1 |
| 102560 | 10 | 2567.8 | 210.3 | 589.7 | 81.8 | 4.4 | 1 |
| 102560 | 50 | 3621.1 | 705.7 | 580.3 | 84.7 | 6.2 | 1 |
| 102560 | 100 | 2567.8 | 210.3 | 589.7 | 81.8 | 4.4 | 1 |
| 102560 | 200 | 4270.0 | 768.7 | 391.3 | 35.7 | 10.9 | 1 |
| 102560 | 500 | 8157.7 | 1898.1 | 412.8 | 45.6 | 19.8 | 1 |
| 111600 | 0 | 1974.0 | 401.8 | 481.5 | 95.1 | 4.1 | 1 |
| 111600 | 10 | 1979.0 | 114.1 | 427.6 | 41.5 | 4.6 | 1 |
| 111600 | 50 | 2512.4 | 458.7 | 442.3 | 47.1 | 5.7 | 1 |
| 111600 | 100 | 3682.8 | 791.7 | 434.7 | 52.7 | 8.5 | 1 |
| 111600 | 200 | 4993.3 | 1289.5 | 460.6 | 63.0 | 10.8 | 1 |
| 111600 | 500 | 15338.3 | 2675.8 | 802.3 | 84.8 | 19.1 | 1 |
| 120640 | 0 | 2667.4 | 466.8 | 690.8 | 156.3 | 3.9 | 1 |
| 120640 | 10 | 3502.8 | 202.2 | 747.5 | 88.4 | 4.7 | 1 |
| 120640 | 50 | 3105.9 | 559.7 | 524.8 | 46.3 | 5.9 | 1 |
| 120640 | 100 | 3805.8 | 326.5 | 561.0 | 54.6 | 6.8 | 1 |
| 120640 | 200 | 5648.2 | 745.5 | 555.3 | 35.4 | 10.2 | 1 |
| 120640 | 500 | 10876.7 | 1011.9 | 618.9 | 85.9 | 17.6 | 1 |
| 129680 | 0 | 3324.1 | 376.3 | 766.6 | 158.5 | 4.3 | 1 |
| 129680 | 10 | 2884.7 | 272.2 | 616.8 | 46.6 | 4.7 | 1 |
| 129680 | 50 | 4415.2 | 1241.0 | 623.0 | 85.9 | 7.1 | 1 |
| 129680 | 100 | 5015.7 | 1087.5 | 601.6 | 50.5 | 8.3 | 1 |
| 129680 | 200 | 10735.2 | 2326.4 | 908.8 | 117.9 | 11.8 | 1 |
| 129680 | 500 | 13058.8 | 2272.6 | 681.5 | 79.2 | 19.2 | 1 |
| 138720 | 0 | 3752.0 | 306.6 | 968.0 | 154.9 | 3.9 | 1 |
| 138720 | 10 | 3452.6 | 150.9 | 693.3 | 58.9 | 5.0 | 1 |
| 138720 | 50 | 4522.5 | 552.6 | 661.6 | 69.1 | 6.8 | 1 |
| 138720 | 100 | 5795.3 | 1111.6 | 681.7 | 70.4 | 8.5 | 1 |
| 138720 | 200 | 10778.0 | 2625.1 | 1069.2 | 129.4 | 10.1 | 1 |
| 138720 | 500 | 15880.2 | 2267.5 | 814.7 | 130.5 | 19.5 | 1 |
| 147760 | 0 | 4781.5 | 311.2 | 944.6 | 152.0 | 5.1 | 1 |
| 147760 | 10 | 4045.2 | 174.5 | 824.7 | 66.1 | 4.9 | 1 |
| 147760 | 50 | 5830.3 | 1191.5 | 845.5 | 114.8 | 6.9 | 1 |
| 147760 | 100 | 10330.9 | 2871.1 | 1164.6 | 99.9 | 8.9 | 1 |
| 147760 | 200 | 8991.0 | 1905.1 | 855.6 | 91.3 | 10.5 | 1 |
| 156800 | 0 | 5279.3 | 1235.1 | 1011.7 | 174.1 | 5.2 | 1 |
| 156800 | 10 | 4496.8 | 288.0 | 979.7 | 80.2 | 4.6 | 1 |
| 156800 | 50 | 5782.3 | 1143.8 | 968.9 | 155.5 | 6.0 | 1 |
| 156800 | 100 | 10325.0 | 1961.7 | 1334.5 | 71.5 | 7.7 | 1 |
| 156800 | 200 | 11101.9 | 1579.7 | 968.5 | 85.7 | 11.5 | 1 |
| 165840 | 0 | 5349.6 | 296.8 | 1196.5 | 142.4 | 4.5 | 1 |
| 165840 | 10 | 5409.9 | 389.8 | 1149.8 | 71.5 | 4.7 | 1 |
| 165840 | 50 | 7394.1 | 2057.1 | 1083.3 | 86.2 | 6.8 | 1 |
| 165840 | 100 | 11390.7 | 2931.3 | 1464.4 | 126.1 | 7.8 | 1 |
| 165840 | 200 | 12595.2 | 1994.5 | 1140.4 | 125.7 | 11.0 | 1 |
| 174880 | 0 | 6231.0 | 1489.3 | 1515.7 | 194.3 | 4.1 | 1 |
| 174880 | 10 | 7394.8 | 415.5 | 1611.3 | 106.2 | 4.6 | 1 |
| 174880 | 50 | 7345.8 | 1510.2 | 1227.3 | 118.5 | 6.0 | 1 |
| 174880 | 100 | 9764.4 | 2724.4 | 1333.5 | 203.9 | 7.3 | 1 |
| 174880 | 200 | 14443.1 | 2256.8 | 1227.7 | 167.4 | 11.8 | 1 |
| 183920 | 0 | 6558.3 | 200.2 | 1674.2 | 328.3 | 3.9 | 1 |
| 183920 | 10 | 7958.2 | 383.8 | 1800.8 | 110.5 | 4.4 | 1 |
| 183920 | 50 | 8371.2 | 1596.2 | 1325.0 | 189.4 | 6.3 | 1 |
| 183920 | 100 | 11133.2 | 3602.7 | 1402.1 | 178.3 | 7.9 | 1 |
| 183920 | 200 | 14618.1 | 2356.3 | 1416.3 | 184.5 | 10.3 | 1 |

**Table 2:** FASTCORE running time improvement due to the preprocessing with FASTERCC. Benchmarks were performed on models with between 10,600 and 93,520 reactions and core sets of 10 to 500 reactions. Preprocessing with FASTERCC shortened the runtime in 48 of 50 reported runs.

| **Input Model Size** | **Number of Core reactions** | **FASTCORE Time (No preprocessing)** | **Time with FASTERCC preprocessing** | **Speedup** |
| --- | --- | --- | --- | --- |
| 10600 | 10 | 4.4 | 3.0 | 1.5 |
| 10600 | 50 | 4.4 | 3.3 | 1.3 |
| 10600 | 100 | 4.5 | 3.5 | 1.3 |
| 10600 | 200 | 5.1 | 3.3 | 1.6 |
| 10600 | 500 | 4.8 | 2.7 | 1.8 |
| 21200 | 10 | 16.3 | 11.6 | 1.4 |
| 21200 | 50 | 16.2 | 12.2 | 1.4 |
| 21200 | 100 | 17.9 | 10.8 | 1.7 |
| 21200 | 200 | 16.6 | 11.4 | 1.5 |
| 21200 | 500 | 16.6 | 11.4 | 1.4 |
| 30240 | 10 | 38.1 | 27.5 | 1.4 |
| 30240 | 50 | 31.7 | 23.9 | 1.3 |
| 30240 | 100 | 35.3 | 24.5 | 1.4 |
| 30240 | 200 | 32.3 | 25.7 | 1.3 |
| 30240 | 500 | 37.9 | 26.0 | 1.5 |
| 39280 | 10 | 52.1 | 41.0 | 1.3 |
| 39280 | 50 | 47.3 | 37.7 | 1.3 |
| 39280 | 100 | 52.6 | 37.4 | 1.4 |
| 39280 | 200 | 43.7 | 37.2 | 1.2 |
| 39280 | 500 | 50.0 | 49.3 | 1.0 |
| 48320 | 10 | 88.3 | 61.4 | 1.4 |
| 48320 | 50 | 108.4 | 66.5 | 1.6 |
| 48320 | 100 | 80.5 | 64.6 | 1.2 |
| 48320 | 200 | 118.2 | 65.4 | 1.8 |
| 48320 | 500 | 100.3 | 71.0 | 1.4 |
| 57360 | 10 | 121.6 | 80.0 | 1.5 |
| 57360 | 50 | 157.7 | 110.8 | 1.4 |
| 57360 | 100 | 132.9 | 130.1 | 1.0 |
| 57360 | 200 | 133.0 | 108.5 | 1.2 |
| 57360 | 500 | 130.2 | 92.5 | 1.4 |
| 66400 | 10 | 193.3 | 150.9 | 1.3 |
| 66400 | 50 | 198.0 | 182.7 | 1.1 |
| 66400 | 100 | 222.2 | 172.1 | 1.3 |
| 66400 | 200 | 192.0 | 172.2 | 1.1 |
| 66400 | 500 | 234.2 | 172.2 | 1.4 |
| 75440 | 10 | 291.4 | 267.3 | 1.1 |
| 75440 | 50 | 261.8 | 230.6 | 1.1 |
| 75440 | 100 | 247.6 | 257.1 | 1.0 |
| 75440 | 200 | 268.5 | 260.5 | 1.0 |
| 75440 | 500 | 254.4 | 222.1 | 1.1 |
| 84480 | 10 | 315.6 | 299.9 | 1.1 |
| 84480 | 50 | 402.0 | 336.0 | 1.2 |
| 84480 | 100 | 327.4 | 307.9 | 1.1 |
| 84480 | 200 | 480.6 | 299.3 | 1.6 |
| 84480 | 500 | 481.5 | 276.0 | 1.7 |
| 93520 | 10 | 456.6 | 342.0 | 1.3 |
| 93520 | 50 | 586.5 | 475.1 | 1.2 |
| 93520 | 100 | 463.1 | 447.3 | 1.0 |
| 93520 | 200 | 404.1 | 452.2 | 0.9 |
| 93520 | 500 | 549.8 | 425.4 | 1.3 |

